# Lactoferrin-Derived Peptide Chimera Induces Caspase-independent Cell Death in Multiple Myeloma

**DOI:** 10.1101/2024.10.24.620137

**Authors:** Young-Saeng Jang, Shima Barati Dehkohneh, Jaewon Lim, Jaehui Kim, Donghwan Ahn, Sun Shim Choi, Seung Goo Kang

## Abstract

Lactoferrin-derived peptide chimera is a synthetic peptide that mimics the functional unit of lactoferrin with antibacterial activity. Antimicrobial peptides are promising candidates for cancer treatment. Although LF is known to have anticancer effects, to the best of our knowledge, its effects on multiple myeloma have not yet been studied. In this study, we explored the potential of lactoferrin-derived chimera for the treatment of multiple myeloma, a malignant clonal plasma cell disease of the bone marrow. Lactoferrin-derived chimera effectively inhibited MM1S, MM1R, and RPMI8226 multiple myeloma cell growth and induced the early and late phases of apoptosis, but not in normal peripheral blood mononuclear cells. Furthermore, lactoferrin-derived chimera modulates the relative expression of genes involved in survival, apoptosis, and mitochondrial dysfunction at the transcriptional level. Mitochondrial analysis revealed that lactoferrin-derived chimera triggered oxidative stress in multiple myeloma cells, leading to the generation of reactive oxygen species and a decline in mitochondrial membrane potential, resulting in mitochondrial dysfunction. Although lactoferrin-derived chimera did not cause caspase-dependent cell death, it induced nuclear translocation of apoptosis-inducing factor and endonuclease G, indicating the initiation of caspase-independent apoptosis. Overall, our data suggest that lactoferrin-derived chimera induces caspase-independent programmed cell death in multiple myeloma cell lines by increasing the nuclear translocation of apoptosis-inducing factor/endonuclease G. Therefore, lactoferrin-derived chimera has potential in multiple myeloma cancer therapies.

## Introduction

Lactoferrin (LF), a versatile glycoprotein enriched in milk and granules of neutrophils, has multiple biological roles, such as immunomodulatory, iron transport, anti-inflammatory, antibacterial, antiviral, and antitumor properties [1, 2]. Initially, LF was shown to have a strong bactericidal function, mainly mediated by two functional domains: lactoferricin and lactoferrampin lacated in the N1 domain of bovine LF [3]. Furthermore, a chimeric form (lactoferrin-derived chimera, LFch) containing a part of lactoferricin (17—30 a.a), and lactoferrampin (265–284 a.a) was created by fusion and surpassed individual peptides in bactericidal activity at low doses and short incubation times [4].

In cancer therapy, LF demonstrates antitumor effects against numerous types of cancer, including breast cancer, cervical cancer, and leukemia [5–7]. The anticancer properties of LF include arresting the cell cycle, damaging the cytoskeleton, inducing apoptosis, and reducing cell migration [8]. Although the observed harm is evident, the underlying mechanisms responsible for these consequences are yet to be fully understood. LF exerts its anticancer effects via two potential mechanisms. It may trigger signaling pathways that can have detrimental effects on cells by binding to proteoglycans, glycosaminoglycans, and sialic acid, which are more abundant in cancer cells than in heathy cells [9]. The other mechanism may involve the disruption of iron metabolism, which is critical for cancer cell survival [10], due to the ability of LF to bind to iron ions [2].

The enhanced interaction of LFch with negatively charged model membranes amplifies the biological activity of these antimicrobial peptides through chimerization, mimicking the native structural orientation of the molecule, which is superior to the native LF form [4]. Therefore, we hypothesize that LFch possesses a strong anticancer effect similar to that of LF and is potentially even superior as it triggers apoptosis in both HepG2 and Jurkat cell lines [11].

In this study, we aimed to explore the antitumor effects of LFch in human multiple myeloma cell lines and clarify the underlying molecular mechanisms. This investigation may provide a theoretical basis for potential future clinical applications in multiple myeloma treatment, which is the second most common hematologic malignancy in adults and the most refractory form of malignancy [12].

## Results

### LF chimera inhibit the growth of MM cell lines

To identify the cytotoxicity of LF and its derivative peptide, (LFch, on MM, MM cell lines (MM1S, MM1R, and RPMI8226) were cultured with LFch, and cell growth was determined by colorimetric assays with various concentration of LFch. The proliferation of all evaluated cell lines was severely decreased by LFch at a concentration of 3 μM and above (Fig. S1). Therefore, we proceeded with a next analysis using the same concentration of LFch. Unlike LF, LFch significantly inhibited the growth of MM1S and RPMI 8226 cells, which was also observed in the drug-resistant MM1R cells (Fig. 1A). LFch also decreased Ki67 expression in these cell lines, suggesting that LFch suppresses the proliferation of MM cells (Fig. 1B). To further characterize the cytotoxicity of LFch in MM cell lines, we examined the effect of LFch on MM cell apoptosis. The annexin V-propidium iodide (PI) double-positive population significantly increased in all three MM cell lines, indicating that LFch induces both early– and late-phase apoptosis, and is pivotal for triggering programmed cell death (PCD) in MM cell lines (Fig. 1C). In contrast, LFch did not induce apoptosis in activated CD19+ B cells in the normal PBMCs (Fig. 1D). Collectively, these results suggest the potential specificity of LFch as a promising therapeutic candidate for MM.

**Fig. 1.**
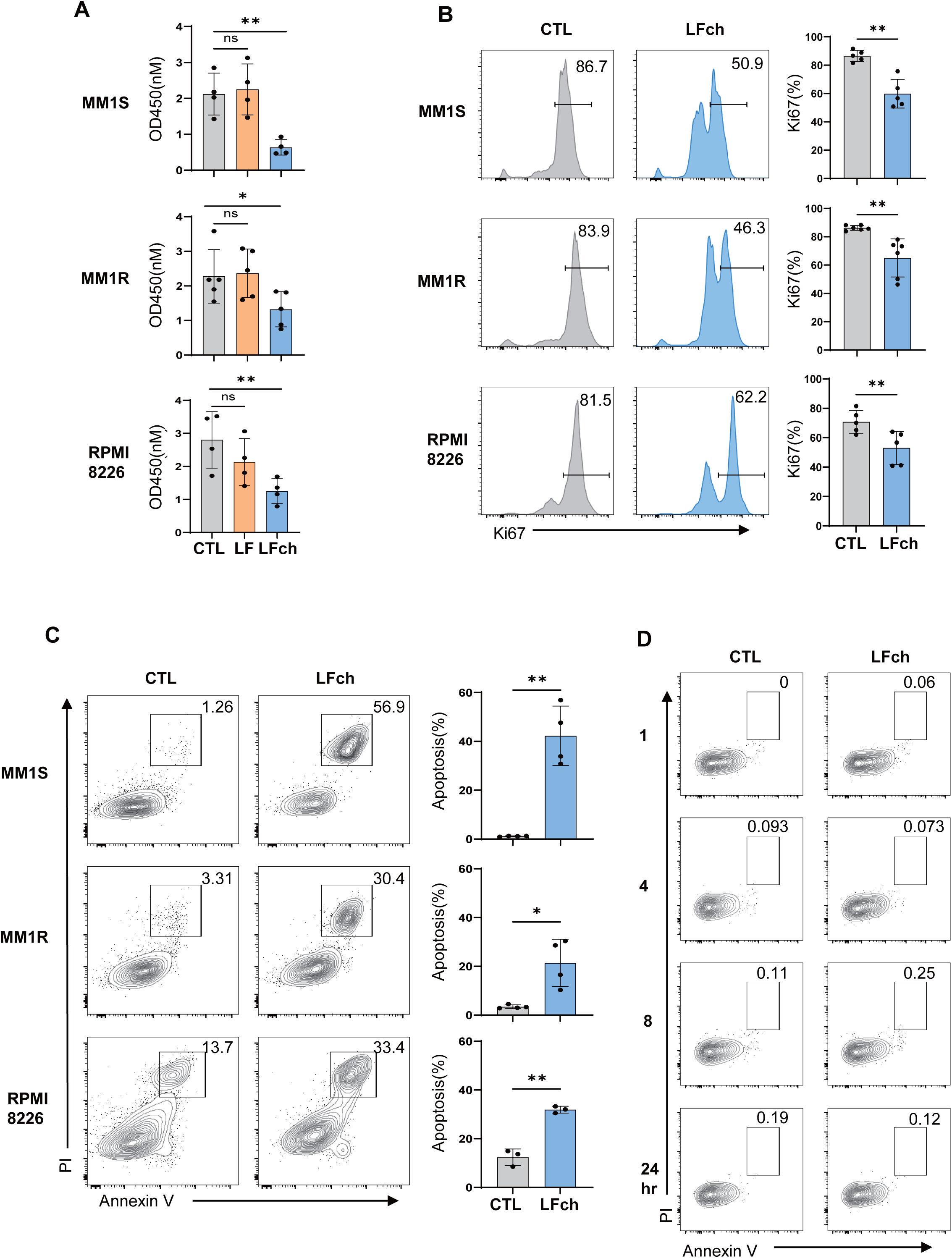
LFch induces cytotoxicity in MM cell lines, but not in normal PBMCs. MM cell lines (MM1S, MM1R, RPMI8226) were treated with 3µM of LF or LFch for 48 h. Growth inhibition by LF or LFch was assessed using the CCK-8 assay (A). Ki67 expression (B) and annexin V^+^PI^+^ apoptotic cells (C) were detected by flow cytometry. (D) Human PBMCs were cultured with 5 µg/ml of anti-human IgM for 48 h, then treated with LFch for indicated times and apoptosis was determined by flow cytometry. Data represent mean plus or minus the standard deviation (± SD) of 3 independent experiments. Statistical analyses were performed using the GraphPad Prism software (GraphPad Software, Inc., La Jolla, CA, USA). (* P < 0.05, ** P < 0.01, versus untreated control). LFch, lactoferrin-derived chimera; MM, multiple myeloma

### Reactive oxygen species (ROS)-induced mitochondrial dysfunction is involved in LFch-induced MM cell apoptosis

To understand the mechanisms underlying the antitumor effects of LFch in MM, we used RNA-sequencing to reveal the genetic landscape. MM cells treated with LFch (3 µM) were subjected to a comprehensive omics study that illuminated potential gene targets and signaling pathways modulated by LFch. RNA sequencing revealed that LFch treatment increased the transcriptional levels of genes related to ROS-induced mitochondrial dysfunction and other cellular responses to cellular stress and DNA damage. In contrast, LFch decreased the expression of cell survival-associated genes in all three MM cell lines (Fig. 2A, B, and C, Fig. S2). Of the 34 common genes identified in MM1S, MM1R, and RPMI8226 cells involved mitochondrial dysfunction and oxidative phosphorylation genes, represented by MT_ND2, MT_ATP6, MT_ND1, MT_ND3, and iron metabolism and oxidative stress genes (*ISCU, FTL,* and *HMOX1*) associated with iron metabolism, revealed the impact of LFch on cellular iron storage, biogenesis of iron-sulfur clusters, and cellular response to oxidative stress (Fig. 2D).

**Fig. 2.**
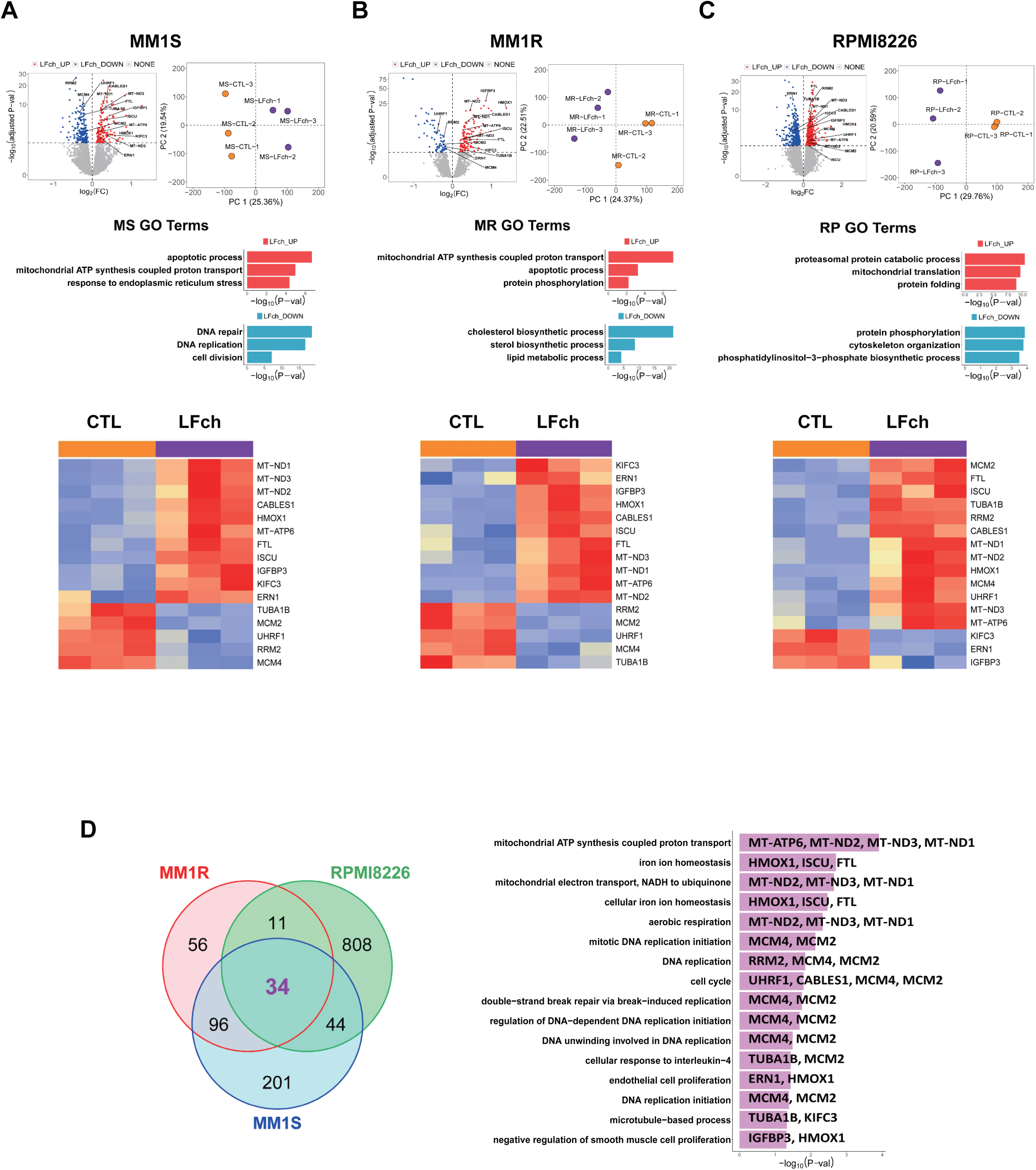
Transcriptomic analyses of LFch-affected MM cells. (A–C) RNA sequencing results depict differentially expressed genes (DEGs) and gene ontology (GO) analysis after LFch (3 µM) treatment on three MM cell lines for 24 h. Volcano plot and Heatmap of genes represent the distribution of DEGs compared to untreated control. Genes labeled in red are significantly upregulated and blue-labeled genes are significantly downregulated. (D) The Venn diagram (left) shows the overlapping number of upregulated gene and 34 common DEGs in three multiple myeloma cell lines. The graph on the right represents the functional classification of 34 common DEGs. LFch, lactoferrin-derived chimera; MM, multiple myeloma

### Mitochondrial dysfunction plays a crucial role in cell death induced by LFch

An examination of LFch-mediated transcriptional changes in MM cell lines revealed the potential of LFch to induce cell death in MM cells through mitochondrial dysfunction. Mitochondrial damage causes the release of several mitochondrial proteins, including ROS, which initiate a PCD cascade[13]. ROS damage cellular components and create a domino effect that disrupts the mitochondrial membrane potential (ΔΨm)[14]. We investigated this complex process by examining the role of ROS in LFch-induced mitochondrial dysfunction. LFch treatment significantly increased ROS levels, which were recovered by the ROS scavenger N-acetylcysteine (NAC) (Fig. 3A). Consequently, cytotoxicity of LFch in MM cell lines was rescued by NAC treatment, highlighting the ROS-dependent mitochondrial destructive effect of LFch (Fig. 3B). Furthermore, LFch treatment resulted in an increase in the TMRE^lo^ fraction of MMs, which is indicative of a reduction in mitochondrial membrane potential (Fig. 3C). Collectively, LFch has the ability to trigger mitochondrial malfunction and programmed cell death (PCD) in MM cell lines.>

**Fig. 3.**
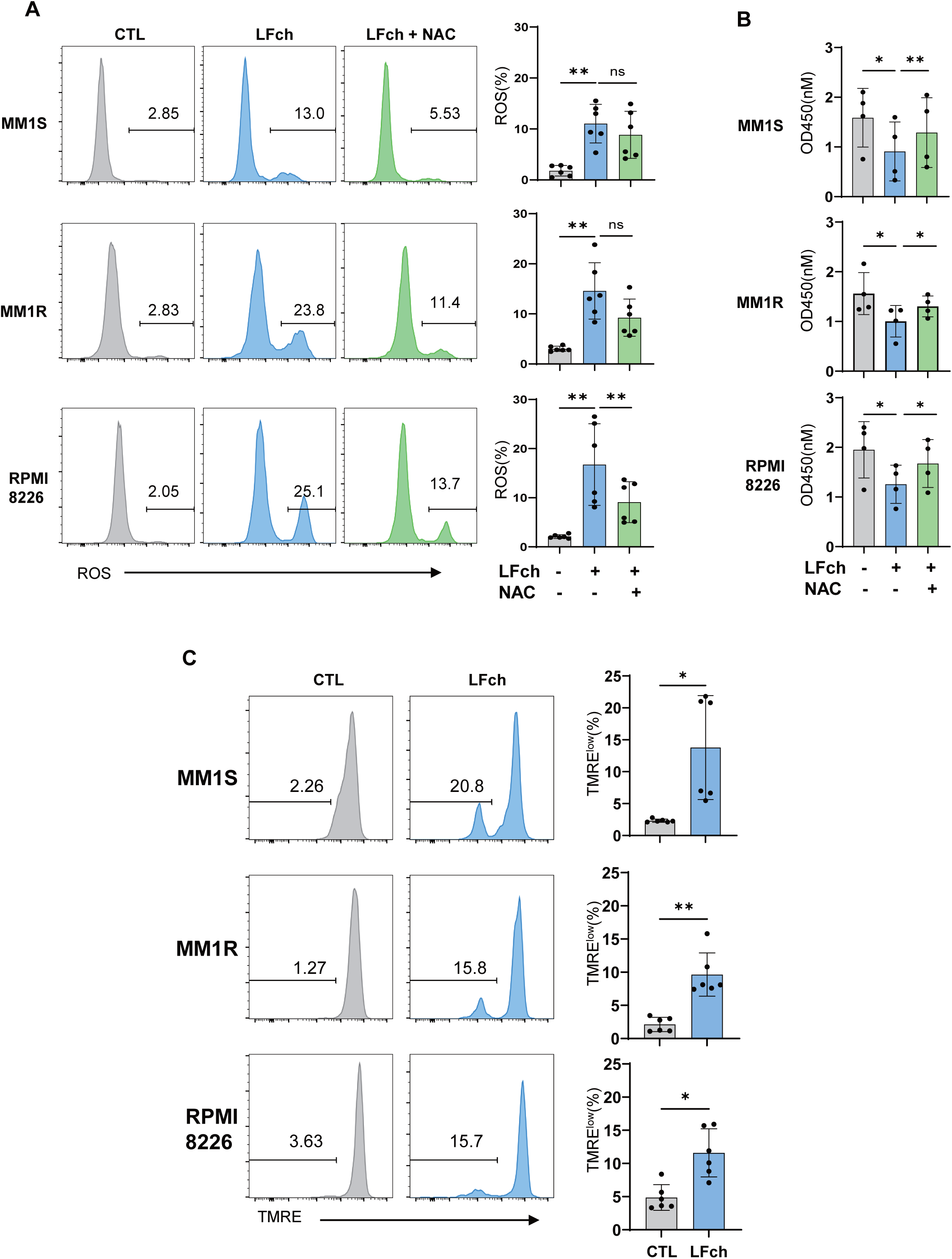
LFch increases the mitochondria ROS and membrane potential in MM cells. (A) MM cells were treated with LFch for 2 h and mitochondria ROS marked by MitoSOX probe was measured using flow cytometry. N-acetylcysteine (NAC, 10 nM) was added to the cultures for ROS scavenging. (B) MM cells were cultured as in A for 48 h, and cell viability was assessed by CCK-8 assay. (C) MM cells were treated with LFch for 2 h and quantitative analysis of mitochondrial membrane potential estimated by TMRE fluorescence. Data represent mean ± SD of three independent experiments (*P < 0.05, **P < 0.01 versus untreated control). LFch, lactoferrin-derived chimera; MM, multiple myeloma

### LFch exerts antitumor activities via caspase-independent apoptosis

Next, to elucidate the mechanism by which LFch affects MM cells, confocal microscopy was performed. We detected penetration of LFch into the cytosol of all MM cell lines (Fig. 4A). This may be because of the general characteristics of LFch as a peptide involved in mitochondrial damage. In the study of PCD pathways, we focused on the role of caspase-3 in LFch-induced apoptosis of MM cells. Notably, western blot analysis revealed the significant absence of caspase-3 expression after LFch treatment. In contrast, the positive control, bortezomib (BTZ), which can induce caspase-3-dependent PCD[15], showed increased expression (Fig. 4B). These results strongly suggest that LFch exerts antitumor effects on MM cells through caspase-3-independent apoptotic pathways. These findings contribute to our understanding of the complex cellular interactions of LFch and further highlight it as a promising candidate for the development of new caspase 3-independent targeted therapies for the treatment of MM. After identifying ROS-induced mitochondrial dysfunction, we sought to elucidate the downstream effects of LFch-induced mitochondrial dysfunction. We investigated the cytoplasmic-to-nuclear translocation of apoptosis-inducing factor (AIF) and endonuclease G (EndoG), which are known to trigger caspase-independent death pathways. We investigated the nuclear translocation of AIF and EndoG using western blotting. In all three MM cell lines, LFch augmented the nuclear translocation of AIF and EndoG (Fig. 4C). This relocation not only increased their movement, but also played a central role in initiating chromatin condensation and DNA fragmentation, which are hallmarks of phase 1 apoptosis. DNA cleavage orchestrated by AIF and EndoG nuclear translocation underscores the role of LFch in caspase-independent death. This mechanism further supports the argument that LFch is a potent inducer of apoptosis in MM cells, providing a unique and promising opportunity for therapeutic interventions. These findings help understand the complex interactions of LFch in the cell and shed light on the specific molecular events that drive its antitumor activity. Identification of AIF and EndoG as key players in this process will provide valuable information for the development of targeted therapies for MM.

**Fig. 4.**
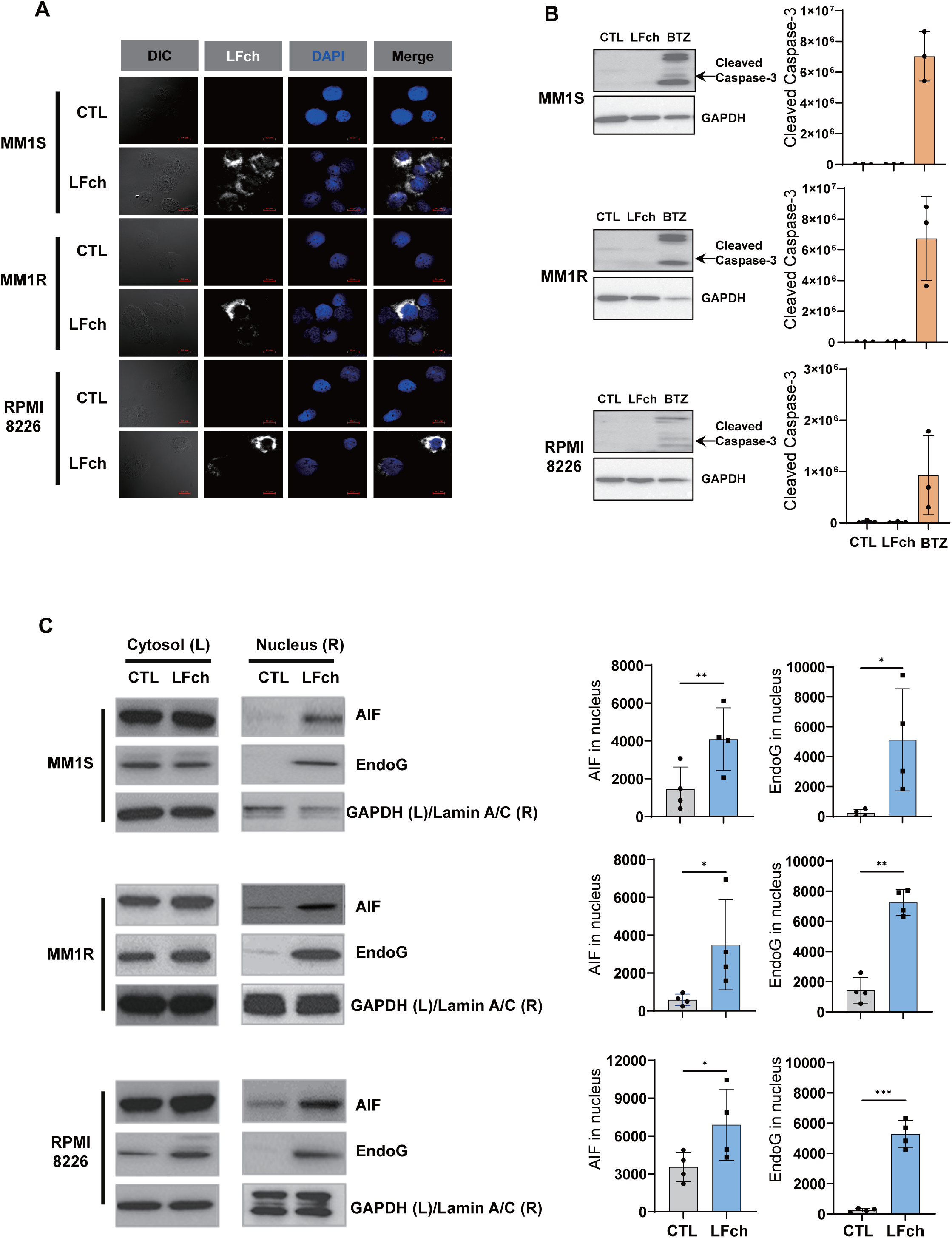
Induction of caspase-independent but AIF and EndoG-dependent apoptosis by LFch. (A) MM cells were treated LFch for 30 min and cellular distribution of LFch was determined by immunocytochemistry. (B) The cleavage of caspase-3 was assessed by immunoblotting in MM cells treated with LFch for 24 h. Bortezomib (BTZ, 25 nM) served as a positive control. (C) The nuclear translocation of AIF and EndoG in cytosol or nuclear fraction of LFch-treated MM cells were determined by Immunoblotting. Data represent mean ± SD of 3 independent experiments (*P < 0.05, **P < 0.01, **P < 0.001 versus untreated control). AIF, apoptosis-inducing factor; EndoG, endonuclease G; LFch, lactoferrin-derived chimera; MM, multiple myeloma

## Discussion

Although treatment approaches for MM have undergone major advancements in recent years, patients with MM frequently experience relapse after one or more treatment regimens or develop drug resistance, along with accompanying side effects and long-term complications. Hence, novel therapeutic approaches are required to address the constraints associated with current treatments [12, 16]. This study elucidates the anticancer characteristics of LFch, which effectively suppresses the proliferation of human MM cell lines, including MM1S, MMIR, and RPMI8226, and triggers PCD, but has no impact on normal human B cells. Additionally, LFch has been shown to have a diverse array of molecular targets that regulate the growth and survival of MMs.

LFch was initially created to mimic the functional components of LF and to function as an antimicrobial peptide (AMPs) [4]. AMPs have attracted considerable interest as promising anticancer drugs because of their distinctive characteristics and mechanisms of action. AMPs possess the crucial characteristic of specifically targeting and disrupting cancer cell membranes, allowing them to function as anticancer drugs [17]. This selectivity is mostly ascribed to disparities in the lipid composition of cancer cell membranes, in contrast to those of normal cells [18]. For example, cancer cells frequently display negatively charged phosphatidylserine on their outer membranes. They can interact with the positively charged nature of many AMPs, causing damage to the cell membrane and resulting in cell death. This process is similar to microbial membrane damage by AMPs, demonstrating a dual function in both antimicrobial and anticancer properties. Furthermore, certain AMPs that containing positively charged and lipid-loving amino acids damage the cell membrane and enhance the passage of hydrophobic mitochondrial membranes, leading to the initiation of apoptosis [19]. Similar to other AMPs that have anticancer effects, LFch also rapidly internalizes into malignant B cells, but not into normal activated B cells, within 30 min. MM is characterized by hypersialylated cell surface membranes [20], and LF preferentially binds to sialic acids [9]. The LF-derived peptide, LFch, may exhibit selective binding to malignant B cells rather than normal B cells, and subsequently penetrate cancer cells. However, the underlying mechanisms require further investigation.

We demonstrated that LFch induces PCD in MM cells without caspase activation. This process occurs by facilitating the translocation of AIF and EndoG from the cytoplasm to the nucleus. Gene profiling of MM cells affected by LFch also demonstrated a notable increase in mitochondrial dysfunction. Induction of apoptosis in tumor cells has been widely acknowledged as a crucial strategy for cancer treatment. Nevertheless, inherent resistance to PCD and swift recurrence of tumors following an initial positive reaction to chemotherapeutic drugs are regarded as the most substantial constraints of treatments that rely on inhibiting cell growth as their main mechanism [21]. Consequently, there is an increasing interest in caspase-independent cell death and the discovery of substances that function largely through the caspase-independent pathway [22]. Understanding this process offers valuable insights into how cells can undergo PCD without the involvement of caspase enzymes, thereby identifying prospective targets for therapeutic interventions in disorders where the regulation of PCD is disturbed. Caspase-independent apoptosis is a crucial process in PCD that depends on mitochondrial malfunction and the activity of pro-apoptotic proteins, such as AIF and EndoG [23, 24]. AIF and EndoG typically reside within the intermembrane gap of mitochondria. After receiving a signal for PCD, they are released into the cytoplasm owing to the breakdown of the mitochondrial membrane, and then move to the nucleus. Within the nucleus, AIF attaches to DNA and triggers the compaction of chromatin as well as the fragmentation of DNA on a wide scale [23]. EndoG can directly cut DNA, resulting in significant DNA fragmentation [24, 25].

Notably, findings not presented in this context indicate that LFch is capable of initiating cellular processes within the nucleus and triggering PCD. Nevertheless, it is essential to recognize that mitochondrial malfunction and the related cell death mechanism may be just one of the multiple mechanisms contributing to LFch-induced cytotoxicity. Additional research is required to evaluate the efficacy of LFch in treating MM utilizing an in vivo tumor model. This limitation remains in our current study.

Collectively, our findings demonstrate that LFch triggers apoptosis in MM cells, including drug-resistant MM1R cells, via a caspase-independent pathway. These findings indicate that LFch has the potential to be a new and focused strategy for overcoming drug resistance and enhancing patient outcomes in MM.

## Methods

### Resources

Comprehensive information on the reagents and resources used in this study is presented in Table 1.

**Table 1.**
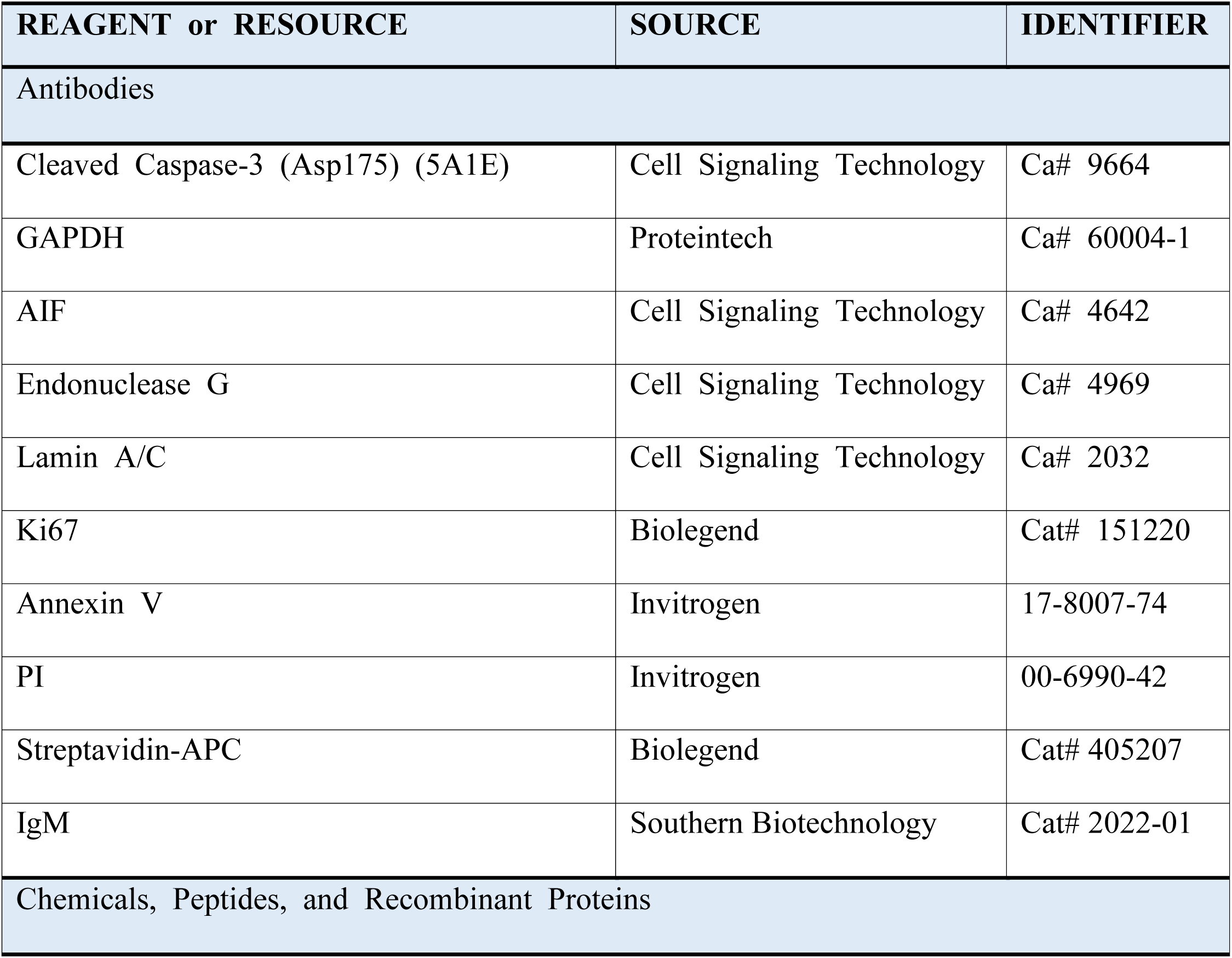
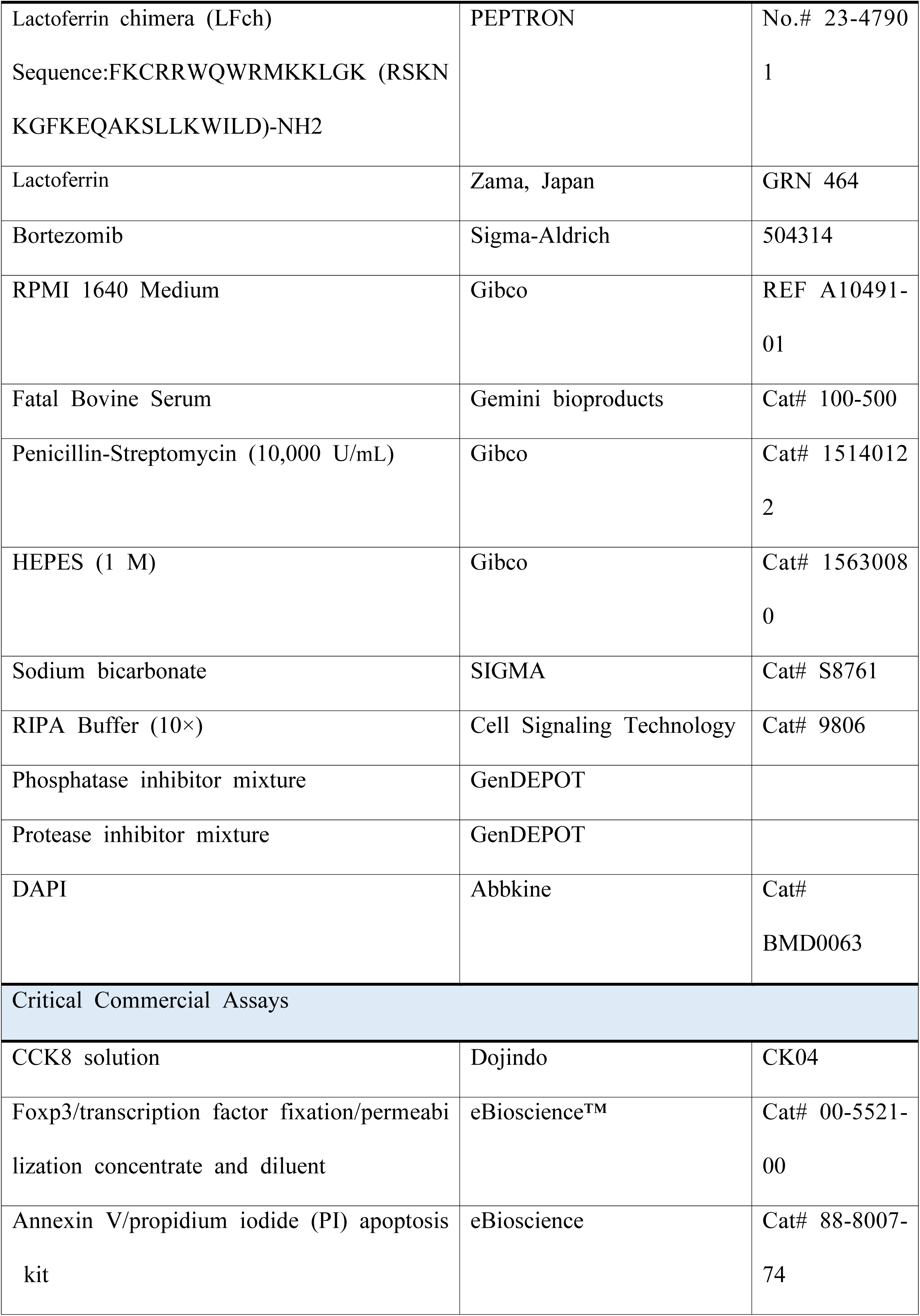
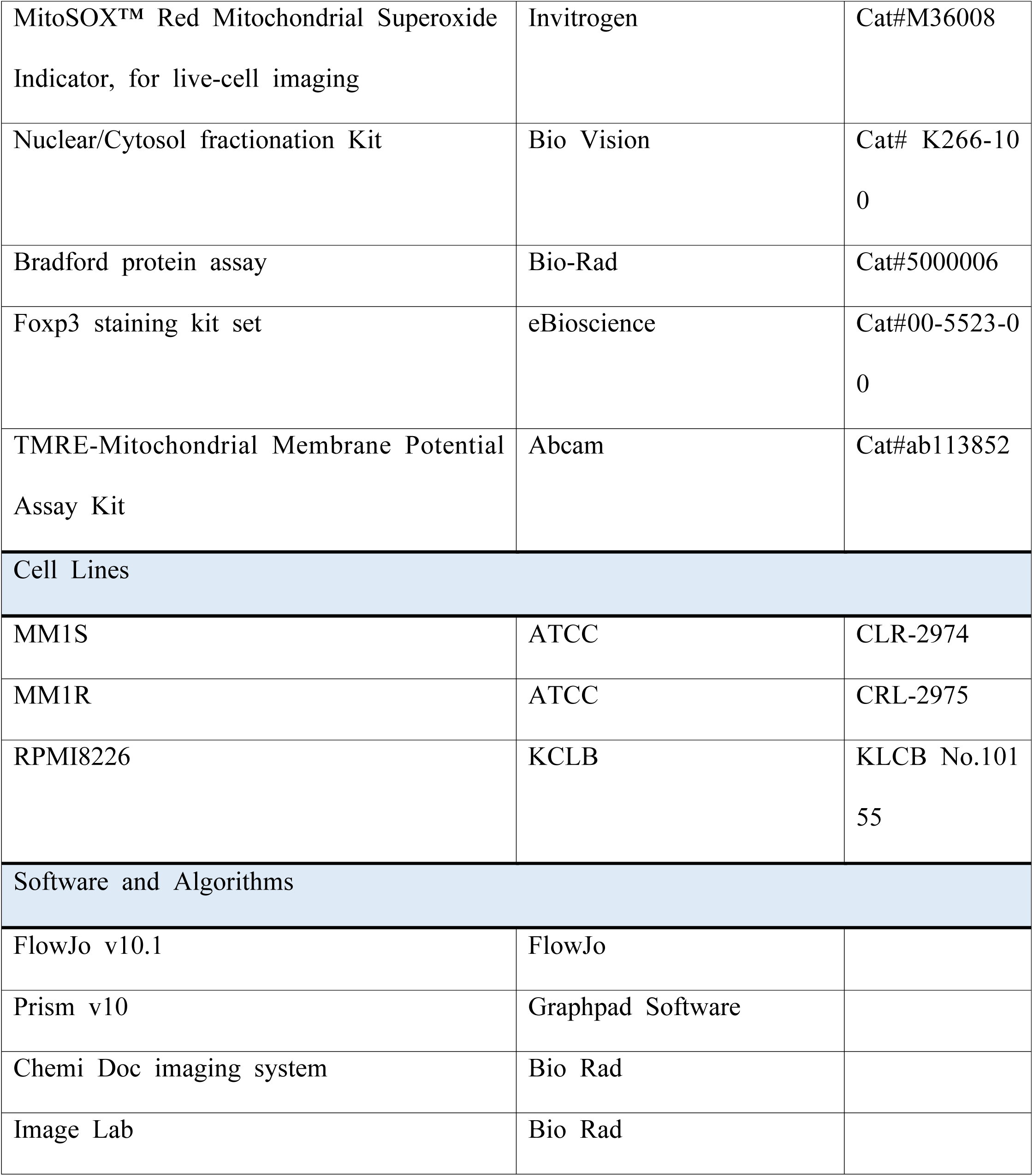
The list of reagents and resources used for this investigation.

| REAGENT or RESOURCE | SOURCE | IDENTIFIER |
| --- | --- | --- |
| Antibodies |  |  |
| Cleaved Caspase-3 (Asp175) (5A1E) | Cell Signaling Technology | Ca# 9664 |
| GAPDH | Proteintech | Ca# 60004-1 |
| AIF | Cell Signaling Technology | Ca# 4642 |
| Endonuclease G | Cell Signaling Technology | Ca# 4969 |
| Lamin A/C | Cell Signaling Technology | Ca# 2032 |
| Ki67 | Biolegend | Cat# 151220 |
| Annexin V | Invitrogen | 17-8007-74 |
| PI | Invitrogen | 00-6990-42 |
| Streptavidin-APC | Biolegend | Cat# 405207 |
| IgM | Southern Biotechnology | Cat# 2022-01 |
| Chemicals, Peptides, and Recombinant Proteins |  |  |
| Lactoferrin chimera (LFch)<br>Sequence:FKCRRWQWRMKKLGK (RSKN<br>KGFKEQAKSLLKWILD)-NH2 | PEPTRON | No.# 23-4790<br>1 |
| Lactoferrin | Zama, Japan | GRN 464 |
| Bortezomib | Sigma-Aldrich | 504314 |
| RPMI 1640 Medium | Gibco | REF A10491-01 |
| Fatal Bovine Serum | Gemini bioproducts | Cat# 100-500 |
| Penicillin-Streptomycin (10,000 U/mL) | Gibco | Cat# 1514012<br>2 |
| HEPES (1 M) | Gibco | Cat# 1563008<br>0 |
| Sodium bicarbonate | SIGMA | Cat# S8761 |
| RIPA Buffer (10×) | Cell Signaling Technology | Cat# 9806 |
| Phosphatase inhibitor mixture | GenDEPOT |  |
| Protease inhibitor mixture | GenDEPOT |  |
| DAPI | Abbkine | Cat#<br>BMD0063 |
| Critical Commercial Assays |  |  |
| CCK8 solution | Dojindo | CK04 |
| Foxp3/transcription factor fixation/permeabi<br>lization concentrate and diluent | eBioscience™ | Cat# 00-5521-00 |
| Annexin V/propidium iodide (PI) apoptosis<br>kit | eBioscience | Cat# 88-8007-74 |
| MitoSOX™ Red Mitochondrial Superoxide Indicator, for live-cell imaging | Invitrogen | Cat#M36008 |
| Nuclear/Cytosol fractionation Kit | Bio Vision | Cat# K266-100 |
| Bradford protein assay | Bio-Rad | Cat#5000006 |
| Foxp3 staining kit set | eBioscience | Cat#00-5523-00 |
| TMRE-Mitochondrial Membrane Potential Assay Kit | Abcam | Cat#ab113852 |
| Cell Lines |  |  |
| MM1S | ATCC | CLR-2974 |
| MM1R | ATCC | CRL-2975 |
| RPMI8226 | KCLB | KLCB No.10155 |
| Software and Algorithms |  |  |
| FlowJo v10.1 | FlowJo |  |
| Prism v10 | Graphpad Software |  |
| Chemi Doc imaging system | Bio Rad |  |
| Image Lab | Bio Rad |  |

### Cell culture

Human MM cell lines, MM1S, MM1R, and RPMI8226, were obtained from the American Type Culture Collection (ATCC, Manassas, VA, USA) and the Korean Cell Line Bank (KCLB, Seoul, Korea). MM cells were maintained in KCLB medium, which consisted of RPMI 1640 medium supplemented with 10% fetal bovine serum (FBS), 1% penicillin/streptomycin, HEPES 1 M, and 7.5% sodium bicarbonate at 37℃ in a humidified atmosphere of 5% CO_2_. Passaging was conducted every 3 days to ensure optimal cell conditions. Human Peripheral blood mononuclear cells (PBMCs) were obtained from Stemcell Technologies (Vancouver, BC, Canada) and cultured in KCLB medium. For activation of primary human B cells, human PBMC were treated with 5 µg/ml of anti-human IgM for 48 h.

### Cell viability and proliferation

The viability of MM cells was measured using a Cell Counting Kit-8 (CCK-8) assay (Dojindo, Japan). Cells (MM1S, MM1R at 1 × 10^5^cells/well; RPMI8226 at 5 × 10^4^ cells/well) were treated with LFch (3µM) for 48 h. Viability was assessed using spectrophotometry at 450 nm. Proliferation analysis was conducted by staining the cells with the Ki67 marker and flow cytometry was used for the analysis.

### Cell apoptosis analysis

MM cells or activated human PBMCs were treated with LFch (3 µM) for 36 h and apoptosis was detected using an Annexin V/PI apoptosis kit (eBioscience) according to the manufacturer’s instructions. Briefly, cells were incubated with Annexin V (1 ml in 20 mL buffer, 1:20) for 20 min at 4℃ and then with PI (1 mL in 40 mL buffer, 1:40) for 5 min in the dark at room temperature.

### Immunoblot analysis

Total protein extraction was performed on cells (MM1S, MM1R, and RPMI8226 at 1 × 10^6^ cells/well) treated with LFch (3 µM). Cells were washed with PBS and lysed in 1× RIPA buffer (Cell Signaling Technology) supplemented with 1× phosphatase inhibitor mixture and 1× protease inhibitor mixture (GenDEPOT) by incubation for 30 min on ice. Nuclear extracts were prepared using a Nuclear/Cytosol Fractionation Kit (BioVision, Mountain View, CA, USA) according to the manufacturer’s instructions. After protein quantification, 20 µg of protein was resolved on NaDodSO4–PAGE gels, transferred onto a polyvinylidene fluoride membrane, and incubated with primary and secondary antibodies. Membranes were scanned using a ChemiDoc imaging system.

### Mitochondrial ROS detection

Changes in mitochondrial ROS were detected using MitoSox Staining. MM cells were treated with LFch (3 µM for 2 h. After washing with PBS, cells were incubated with 1 µM MitoSox solution and analyzed using a Flow Cytometer.

### Mitochondrial membrane potential

Disruption of mitochondrial membrane potential was indicated using TMRE Staining. MM cells were treated with LFch (3 µM) for 2 h. After washing with PBS, the cells were incubated with 200 nM TMRE solution and analyzed using a Flow Cytometer.

### RNA sequencing

MM cells were treated with LFch (3 µM) for 24 h. Total RNA was extracted using the RNAeasy Kit (QIAGEN) according to the manufacturer’s instructions. The RNA library preparation and sequencing reactions were conducted according to standard Illumina protocols by Azenta Life Science (Suzhou, China). Sequencing libraries were generated using the Next® Ultra RNA Library Prep Kit for Illumina® (NEB) following the manufacturer’s protocol.

### Data production and differentially expressed gene identification

Triplicate RNA-seq data were generated from each of the three cell lines, MM1S, MM1R, and RPMI8226, under both control and LF-treated (LFch) conditions. After completing quality checks on the raw sequence data using FastQC (ver. 0.11.9), adapter sequences were trimmed using Trimmomatic (ver. 0.39). The cleaned read sequences were aligned to the reference genome (GRCh38/hg38) using the STAR (ver. 2.7.10a), and HTseq (ver. 0.11.2) was used to count aligned sequencing reads. Differentially expressed genes (DEGs) were identified using normalized read count data and the DESeq2 package in R (ver. 1.38.3) by comparing the control and LFch groups for each of the three cell lines. The two thresholds used for DEG identification were |log2(fold change)| ≥ 0 and adjusted P-value ≤ 0.01.

### Data analysis

Gene Ontology (GO) analysis was performed using DAVID (https://david.ncifcrf.gov/home.jsp), with terms exhibiting a P-value ≤ 0.05 being selected. All other data analyses were performed using R package (ver. 4.2.3). Specifically, principal component analysis (PCA) was conducted using the ‘factoMineR’ package (ver. 2.9), and heatmaps were constructed with z-scores of the normalized read count data using the ‘pheatmap’ package (ver. 1.0.12). Venn diagram analysis was conducted with the ‘VennDiagram’ package (ver. 1.7.3). The ‘ggplot2’ (ver. 3.4.2) and ‘ggrepel’ (ver. 0.9.4) packages in R were used to visualize volcano, PCA, and GO term bar plots.

### Confocal microscopy

The MM cells were incubated with LFch-biot for 1 h, and fixed with 4% paraformaldehyde. The cells were blocked and permeabilized with DPBS containing 10% goat normal serum and 0.1% Triton X-100. Cells were cytospun, and the slides were incubated with SAV-APC (BioLegend, San Diego, CA, USA) and stained with DAPI (Abbkine, Georgia, USA) for nuclear staining. The relative distribution of the fluorochromes was visualized using a supersensitive high-resolution confocal laser scanning microscope LSM880 with Airyscan (Carl Zeiss, Oberkochen, Germany).

### Statistical analysis

Data were analyzed using GraphPad Prism 10 (GraphPad Software, Inc., USA). Statistical analyses were performed using GraphPad Prism 10. Comparisons were performed using two-tailed Student’s t-test or two-way ANOVA. A P-value of ≤ 0.05 was considered statistically significant.

### Data code and availability

The GEO accession number for the RNA-sequencing data reported in this paper is GSE273287.

## Acknowledgments

This research was supported by National Research Foundation of Korea (NRF) grants funded by the Korean government (MSIT) (2021R1A2C1012421 to S. G. K., RS-2024-00341909 to S.S.C., and 2022R1I1A1A01053235 to Y. S. J.).

## Author Contributions

Y.-S.J. and S.G.K. conceived and designed the study. S. B. D., Y.-S. J., J.L., and J.K. performed and analyzed the experiments under the supervision of S.G.K. D.A. performed bioinformatics analysis under the supervision of S.S.C. Y.-S.J., S.B.D., and S.G.K. wrote the manuscript with input from the other authors. All the authors contributed to the manuscript and approved the submitted version.

## Abbreviations

LF: Lactoferrin
LFch: LF chimera
MM: multiple myeloma
ROS: reactive oxygen species
AMPs: antimicrobial peptides
AIF: apoptosis-inducing factor
Endo G: Endonuclease G

## Supporting information

**Fig S1.** MM cells were treated with indicated dose of LFch for 48 h. Growth inhibition of LFch was assessed by CCK-8 assay. LFch, lactoferrin-derived chimera; MM, multiple myeloma.

**Fig S2.** Heatmaps shows the genes associated with apoptosis, mitochondrial dysfunction, oxidative phosphorylation and cell survival in LFch-treated MM cells. LFch, lactoferrin-derived chimera; MM, multiple myeloma.

## References

1. Kowalczyk, P., et al., The Lactoferrin Phenomenon-A Miracle Molecule. Molecules, 2022. 27(9).

2. Baker, H.M. and E.N. Baker, Lactoferrin and iron: structural and dynamic aspects of binding and release. Biometals, 2004. 17(3): p. 209–16.

3. Vogel, H.J., et al., Towards a structure-function analysis of bovine lactoferricin and related tryptophan– and arginine-containing peptides. Biochem Cell Biol, 2002. 80(1): p. 49–63.

4. Bolscher, J.G., et al., Bactericidal activity of LFchimera is stronger and less sensitive to ionic strength than its constituent lactoferricin and lactoferrampin peptides. Biochimie, 2009. 91(1): p. 123–32.

5. Mahidhara, G., et al., Oral administration of iron-saturated bovine lactoferrin-loaded ceramic nanocapsules for breast cancer therapy and influence on iron and calcium metabolism. Int J Nanomedicine, 2015. 10: p. 4081–98.

6. Luzi, C., et al., Apoptotic effects of bovine apo-lactoferrin on HeLa tumor cells. Cell Biochem Funct, 2017. 35(1): p. 33–41.

7. Lu, Y., et al., PFR peptide, one of the antimicrobial peptides identified from the derivatives of lactoferrin, induces necrosis in leukemia cells. Sci Rep, 2016. 6: p. 20823.

8. Zhang, Y., C.F. Lima, and L.R. Rodrigues, Anticancer effects of lactoferrin: underlying mechanisms and future trends in cancer therapy. Nutr Rev, 2014. 72(12): p. 763–73.

9. Cutone, A., et al., Lactoferrin’s Anti-Cancer Properties: Safety, Selectivity, and Wide Range of Action. Biomolecules, 2020. 10(3).

10. Kazan, H.H., C. Urfali-Mamatoglu, and U. Gunduz, Iron metabolism and drug resistance in cancer. Biometals, 2017. 30(5): p. 629–641.

11. Arredondo-Beltran, I.G., et al., Antitumor activity of bovine lactoferrin and its derived peptides against HepG2 liver cancer cells and Jurkat leukemia cells. Biometals, 2023. 36(3): p. 639–655.

12. Malard, F., et al., Multiple myeloma. Nat Rev Dis Primers, 2024. 10(1): p. 45.

13. Perillo, B., et al., ROS in cancer therapy: the bright side of the moon. Exp Mol Med, 2020. 52(2): p. 192–203.

14. Suski, J.M., et al., Relation between mitochondrial membrane potential and ROS formation. Methods Mol Biol, 2012. 810: p. 183–205.

15. Lohberger, B., et al., The Proteasome Inhibitor Bortezomib Affects Chondrosarcoma Cells via the Mitochondria-Caspase Dependent Pathway and Enhances Death Receptor Expression and Autophagy. PLoS One, 2016. 11(12): p. e0168193.

16. Elbezanti, W.O., et al., Past, Present, and a Glance into the Future of Multiple Myeloma Treatment. Pharmaceuticals (Basel), 2023. 16(3).

17. Tornesello, A.L., et al., Antimicrobial Peptides as Anticancer Agents: Functional Properties and Biological Activities. Molecules, 2020. 25(12).

18. Preta, G., New Insights Into Targeting Membrane Lipids for Cancer Therapy. Front Cell Dev Biol, 2020. 8: p. 571237.

19. Hwang, B., et al., The antimicrobial peptide, psacotheasin induces reactive oxygen species and triggers apoptosis in Candida albicans. Biochem Biophys Res Commun, 2011. 405(2): p. 267–71.

20. Daly, J., et al., Targeting hypersialylation in multiple myeloma represents a novel approach to enhance NK cell-mediated tumor responses. Blood Adv, 2022. 6(11): p. 3352–3366.

21. Mohammad, R.M., et al., Broad targeting of resistance to apoptosis in cancer. Semin Cancer Biol, 2015. 35 **Suppl**(0): p. S78–S103.

22. Bhadra, K., A Mini Review on Molecules Inducing Caspase-Independent Cell Death: A New Route to Cancer Therapy. Molecules, 2022. 27(19).

23. Sevrioukova, I.F., Apoptosis-inducing factor: structure, function, and redox regulation. Antioxid Redox Signal, 2011. 14(12): p. 2545–79.

24. Zhdanov, D.D., et al., Regulation of Apoptotic Endonucleases by EndoG. DNA Cell Biol, 2015. 34(5): p. 316–26.

25. Cregan, S.P., et al., Apoptosis-inducing factor is involved in the regulation of caspase-independent neuronal cell death. J Cell Biol, 2002. 158(3): p. 507–17.

