## Supplementary figures and images for "Lactoferrin-Derived Peptide Chimera Induces Caspase-independent Cell Death in Multiple Myeloma"

### supplementary figures 1 and 2

#### Supplement figure 1

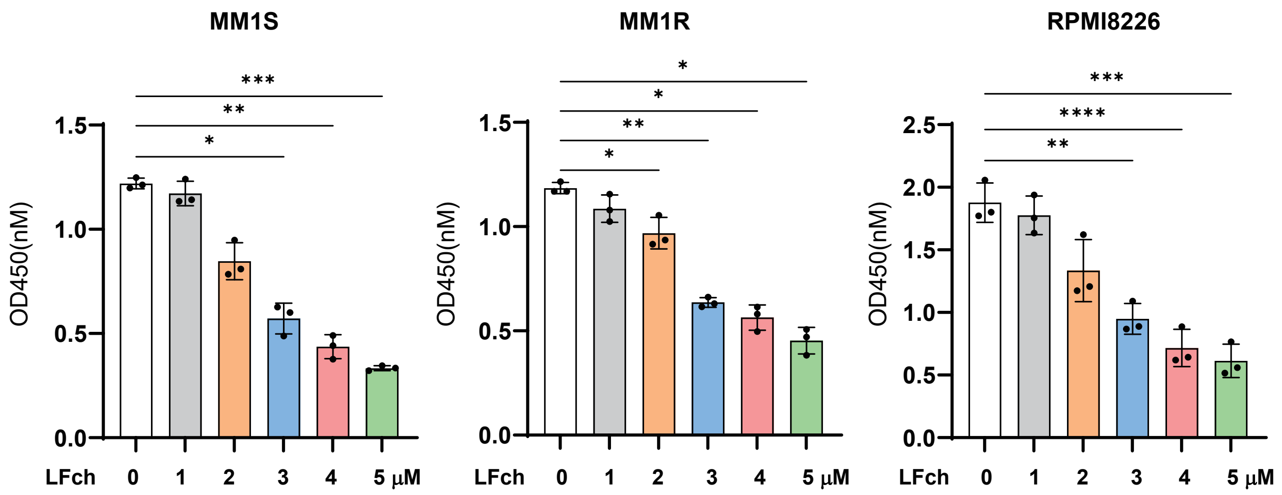

#### Supplement figure 2

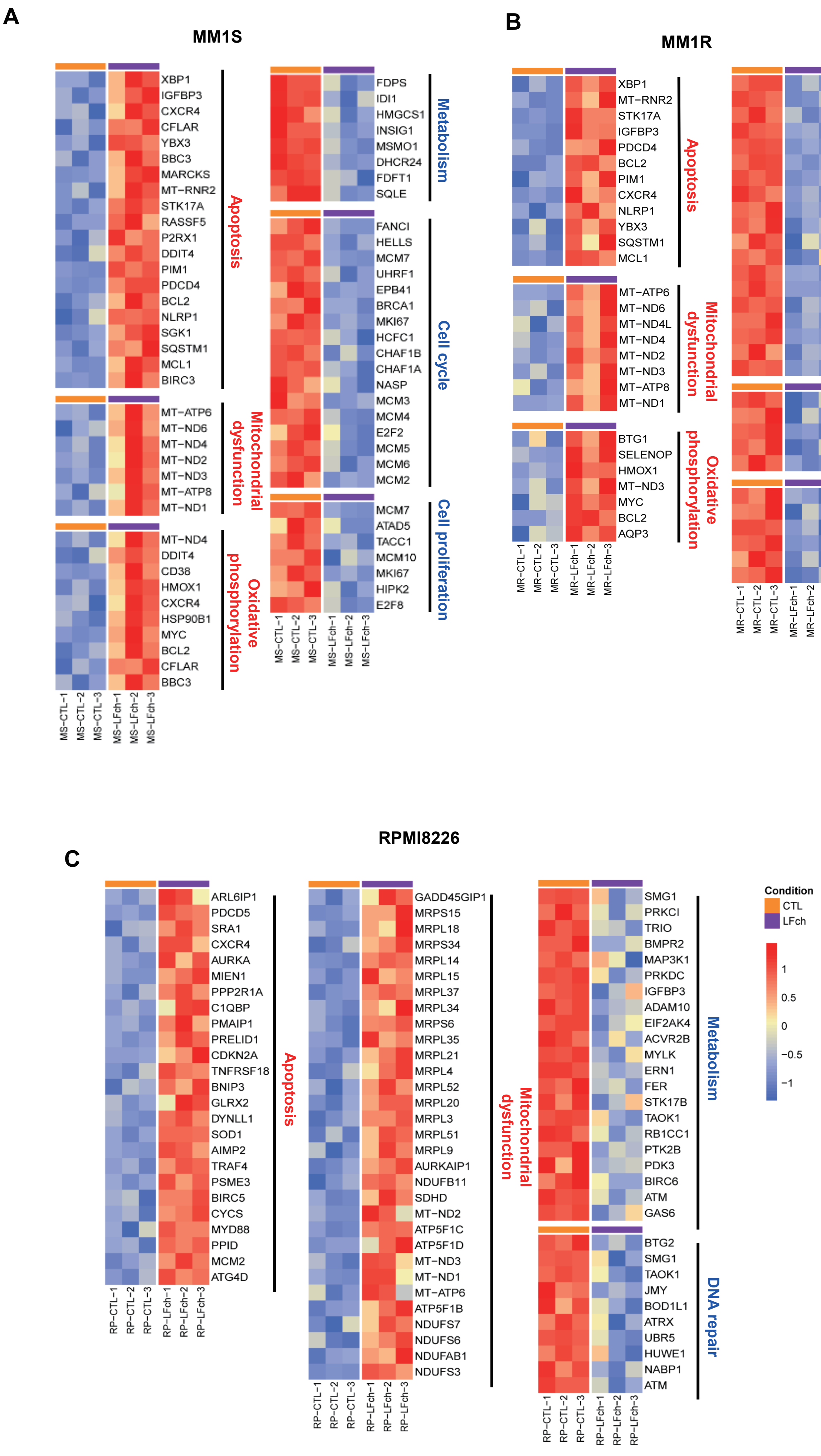
